# Efficiently, accurately, intelligently dissecting bamboo: characteristic of vascular bundle in *Phyllostachys*

**DOI:** 10.1101/2021.12.01.470805

**Authors:** Haocheng Xu, Ying Zhang, Jiajun Wang, Tuhua Zhong, Xinxin Ma, Jing Li, Hankun Wang

**Affiliations:** Institute of New Bamboo and Rattan Based Biomaterials, International Center for Bamboo and Rattan; Key Laboratory of National Forestry and Grassland Administration/Beijing for Bamboo & Rattan Science and Technology, Beijing, 100102, China; Research Institute of Wood Industry, Chinese Academy of Forestry; Key Laboratory of National Forestry and Grassland Administration for Wood Science and Technology, Beijing, 100091, China

**Keywords:** *Phyllostachys*, vascular bundle, fiber volume fraction, radial distribution

## Abstract

A comprehensive understanding of vascular bundles is the key to elucidate the excellent intrinsic mechanical properties of bamboo. This research aims to investigate the gradient distribution of fiber volume fraction and the gradient changes in the shape of vascular bundles along the radial axis in *Phyllostachys*. We constructed a universal transfer-learning-based vascular bundle detection model with high precision of up to 96.97%, which can help to acquire the characteristics of vascular bundles quickly and accurately. The total number of vascular bundles, total fiber sheath area, the length, width and area of fiber sheath of individual vascular bundles within the entire cross-section were counted, and the results showed that these parameters had a strongly positive linear correlation with the outer circumference and wall thickness of bamboo culms, but the fiber volume fraction (around 25.5 %) and the length-to-width ratio of the vascular bundles (around 1.226) were relatively constant. Furthermore, we layered the cross section of bamboo according to the wall thickness finely and counted the characteristics of vascular bundle in each layer. The results showed that the radial distribution of fiber volume fraction decreased exponentially, the radial distribution of the length-to-width ratio of vascular bundle decreased quadratically, the radial distribution of the width of vascular bundle increased linearly. The trends of the gradient change in vascular bundle’s characteristics were found highly consistent among 29 bamboo species in *Phyllostachys*.

**One sentence summary:** A universal vascular bundle detection model can efficiently dissect vascular bundles in *Phyllostachys*, and the radial gradient change of vascular bundles in cross-section are found highly consistent.

## 1. Introduction

Bamboos are monocotyledons and belong to Bambusoideae, Poaceae, and Poales (Clark et al., 2015). Bamboos grow so quickly that they can reach up to 20-30 m tall within 40-50 days. The growth height can reach up to 1-1.5 m per day. Bamboos produce high yields of raw materials for use, as biomass rises by 10-30 percent annually versus 2-5 percent for trees. And once bamboos are planted, they can have sustainable use for years. Furthermore, bamboos require simple processing techniques and are obtained from renewable and ecological resources (Ghavami et al., 2003). Because of its wide range of applications in construction (Sá Ribeiro et al., 2004; Lo et al., 208; Chele et al., 2012; Shi et al., 2019), the development of bamboo resources arouses great attention around the world, especially in tropical and sub-tropical countries with rapidly developing economies (Ruy et al., 2017). The properties of bamboo originate from the composition. The organizational structure of bamboo is mainly composed of vascular bundles and parenchyma (Obataya et al., 2007). The vascular bundles are the reinforcement and transfusion tissues of bamboo, consisting the primary phloem, the fascicular cambium, and the primary xylem. The vascular bundles provide the bamboo with rigidity by stretching continuously in the longitudinal direction of each internode. In contrast, parenchyma, which consist of thin-walled cells, can pass loads and take the role of a composite matrix but contribute little to the rigidity of bamboo. Thus, the mechanical properties of bamboo depend on variation in the morphology and distribution of the vascular bundles in the cross section (Amada & Untao, 2001; Ghavami et al., 2003; Shao et al., 2009; Palombini et al., 2020).

In light of the earlier studies, the morphology and distribution of vascular bundles differed not only among different bamboo species, but also in different locations of the same bamboo species. The results indicate that from base to top of bamboo culm, the number of vascular bundles decreased along the longitudinal direction, while the distribution density increases; in the radial direction, the number of vascular bundles decreases from the outer skin to the inner skin of bamboo culm (Grosser & Liese, 1971; Li et al., 2021). Nakato (1959) divided the culm wall into four equal layers and examined vascular bundles cross-section in each layer. He found no clear relationship between the height from the ground of the internodes and the length-to-width ratio of vascular bundles and that the ratio decreased from the outer to inner layers. Besides, the cross-sectional area of the individual vascular bundle in the outer skin is smaller than that in the inner skin of bamboo culm, whereas the fiber content is higher (Shang et al., 2011). The morphology and distribution density of vascular bundles change continuously from outer skin to inner skin of bamboo culm in the radial direction (Shang et al., 2011; Huang, 2012); the distribution density of vascular bundles in the outer skin is 8 per mm^2^, whereas 2 per mm^2^ in the inner skin (Ray et al., 2005). Generally, the morphology of vascular bundles changes little with bamboo age (Ximena L, 2002). However, the results above are mainly obtained from small-sized samples (Legland et al., 2014; Zhang et al., 2017). Obviously, the radial distribution of fiber volume fraction decreased from outer to inner, while there were three mainstream views on the form of decrease in academia. Some researchers believed that the decrease was linearly (Xian et al., 1990; Li et al., 2011), some researchers proposed the decrease was quadratically (Nogata & Takahashi, 1995; Amada et al., 1997; Ghavami et al., 2003; Sato et al., 2017), and others considered the decrease was exponentially (Lee et al., 2014; Chen, 2019; Xu et al., 2021), which may be related to the species, ages, defects and growth environments of bamboo.

Although the characteristics of vascular bundle in cross section have been explored intensively and extensively, it was still a big challenge to quickly and accurately detect the number and the area of vascular bundles in large quantities and large-sized samples (Legland et al., 2014; Zhang et al., 2017). Machine learning might be one of the most potential approaches to solve this issue. Our research group (Li et al., 2021) constructed a vascular bundle detection model based on YOLO algorithm. It could be used to identify vascular bundles of Moso bamboo and measure the shape of vascular bundles quantitatively. We labeled over 20,000 vascular bundle images to build the detection model, which was a very time-consuming and laborious task. In this research, we applied the former vascular bundle detection model as the source domain. Based on transfer learning (a machine learning method for training models with similar features), only a small number of labeled vascular bundles (1/10 of the source domain) were required for constructing the universal detection model, which could accomplish the detection, positioning, counting and measurement of vascular bundles automatically and simultaneously.

*Phyllostachys* belongs to the Arundinarieae family of Poaceae, Bambusoideae subfamily. There are 50 species of *Phyllostachys* and the distribution center and origin of *Phyllostachys* are in China, most of which have high economic, ecological, medicinal and ornamental value. The planting area of *Phyllostachys* accounts for about 3/4 of the total area in the world, which indicated that *Phyllostachys* is the main management object of bamboo forest in China.

In this study, the universal detection model of vascular bundle was constructed and used to analyze twenty-nine kinds of *Phyllostachys* in cross section, obtain (1) the total number of vascular bundles, (2) the distribution density of vascular bundles, (3) the total area of fiber sheath, (4) the fiber volume fraction, (5) the mean value of single fiber sheath area, (6) the mean value of the length of vascular bundle, (7) the mean value of the width of the vascular bundle and (8) the mean value of the length-to-width ratio of vascular bundle. In addition, the relationship between the characteristics of vascular bundle and the outer circumference and the wall thickness of bamboo was also discussed. Furthermore, the radial changes in fiber volume fraction and the shape (the length, the width and the length-to-width ratio) of vascular bundle were studied. It was expected to guide the modern processing, production and innovative design of bamboo, and to give full play to the potential value of gradient structure.

## 2. Results and discussions

### 2.1 Model validation

The universal detection model was used to detect the vascular bundles in scanned images of twenty-nine kinds of *Phyllostachys* to assess the performance of the detection model. Meanwhile, the number of vascular bundles was also counted manually to check the accuracy of the universal detection model. Figure 1 shows the result for samples detected by the model. The predicted value was the number of vascular bundles identified by the detection model. The false value was the sum of the “missing value” caused by unidentified vascular bundles (especially the vascular bundles located in the outer skin of bamboo which were narrow in shape and small in size) and the “error value” caused by the identification of noise in the background such as dust like vascular bundles. The Precision rate was calculated according to the Equations below:

**Figure 1.**
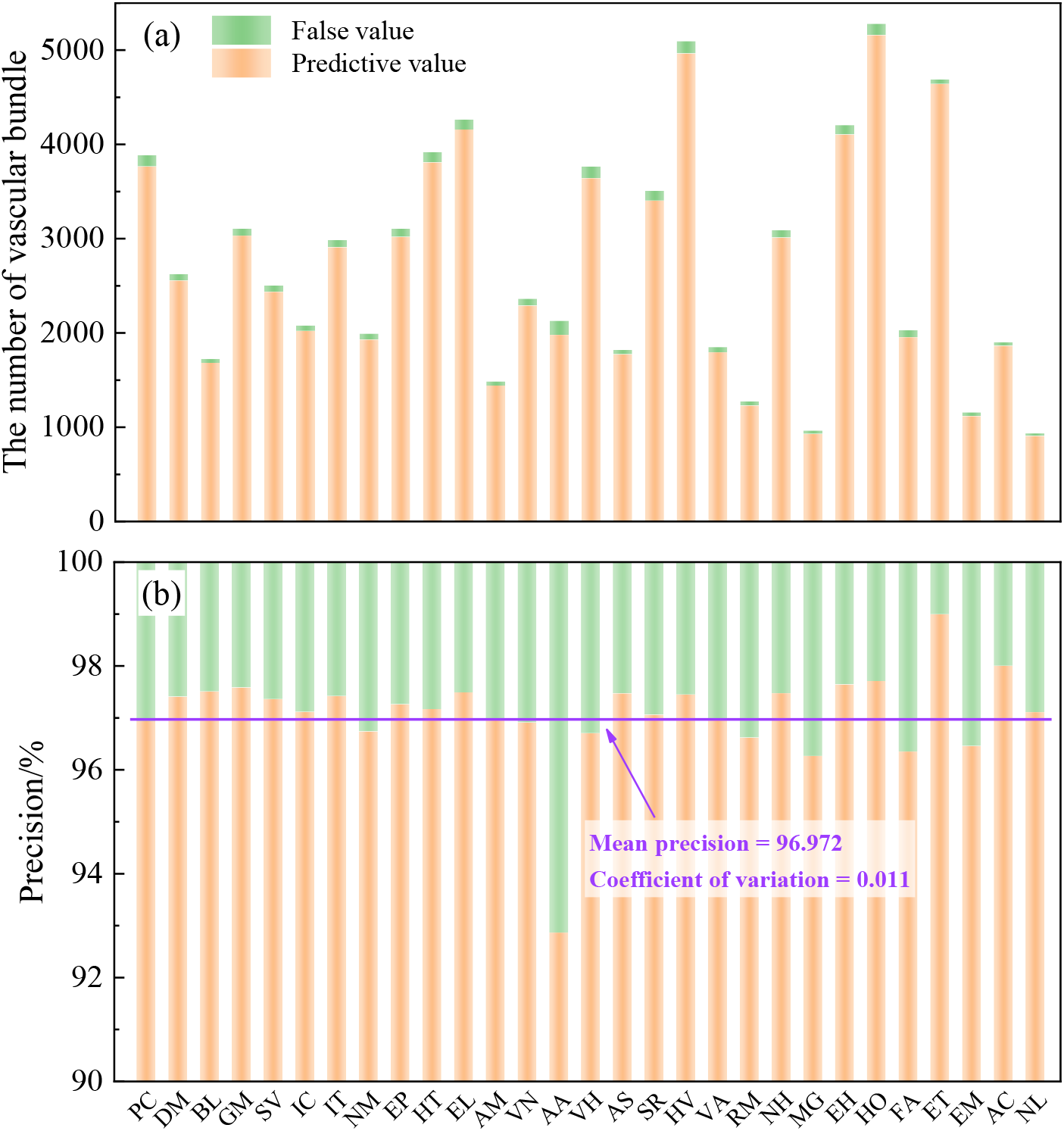
The result for samples detected by the model: **a** vascular bundle identification of bamboo ring; **b** the accuracy of the universal vascular bundle detection model

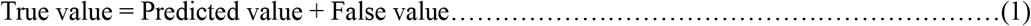

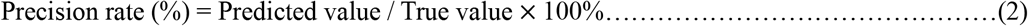

As shown in Figure 1, the accuracy of the universal vascular bundle detection model was high and the precise rate can reach 96.97%. This indicated that the detection model constructed through transfer learning was successful and could perform fast and reliable detection of vascular bundles of *Phyllostachys*. However, unlike the high detection accuracy of the rest of 28 bamboo species, the precision rate of **AA** was relatively low at around 93%, which might be largely due to the light color of the vascular bundles that have the light contrast with the color of the parenchyma cells in the cross-section of **AA**. In machine learning, we attempted to construct a model in which all weights are relatively balanced to prevent over-fitting. In this study, if the final accuracy of the model needs to be improved, it was advisable to increase the labelled volume of the vascular bundles of **AA** to increase the weights of the model.

### 2.2 The vascular bundle characteristics in the cross-section

The universal detection model was used to predict the cross-sectional image of twenty-nine kinds of *Phyllostachys*, the characteristics of vascular bundle were obtained in Table 1. The fiber volume fraction was calculated to be 17.69-32.65%, with an average of 25.50 ± 3.51%; and the mean value of the length-to-width ratio of the vascular bundle was 1.037-1.401, with an average of 1.226 ± 0.091. Whereas, there were apparent differences in the other characteristics of vascular bundle of different kinds of *Phyllostachys*. It possibly results from the outer circumference and the wall thickness of a bamboo culm.

**Table 1.**
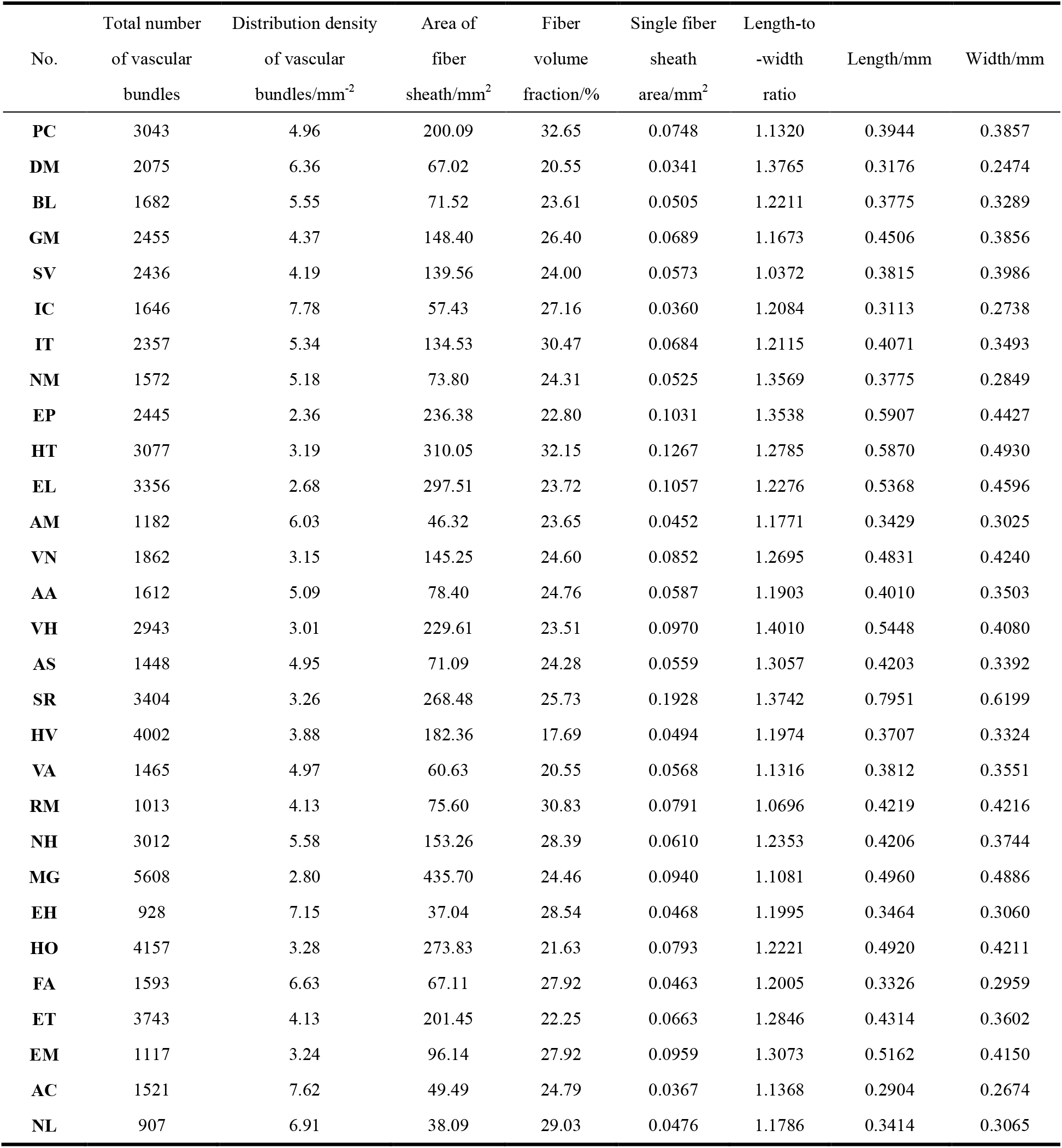
Vascular bundle information in cross section

Table 2 shows the Pearson correlation analysis of the outer circumference and the wall thickness of the bamboo ring and the characteristics of vascular bundle. There was a positive correlation between the total number of vascular bundles, the total area of fiber sheath, the mean value of single fiber sheath area, the mean value of the length of vascular bundle, the mean value of the width of the vascular bundle and the outer circumference and the wall thickness of bamboo (at the 0.01 level, bilateral). Among them, the correlation between the total number of vascular bundles and the total area of fiber sheath and the outer circumference and the wall thickness of the bamboo ring was the hightest. The factors that jointly determine the total number of vascular bundles and the total area of fiber sheath was the size of the cross section. The larger the cross section of the bamboo ring, the more vascular bundles it contains and the larger the total area of fiber sheath it has. Therefore, the two have an obvious positive correlation, and the size of the cross section was determined by the outer circumference and the wall thickness of bamboo. Meanwhile, there was no significant difference in the fiber volume fraction in the cross section among different species, it was expected that *Phyllostachys* with larger outer circumference and wall thickness have increased the length and width of a single vascular bundle. Conversely, the distribution density of vascular bundles was significantly negatively correlated with the outer circumference and the wall thickness of bamboo (0.01 level, bilateral).

**Table 2.** Pearson correlation analysis of the macro characteristics and the characteristics of vascular bundle

| Characteristic of bamboo ring | Outer circumference /mm | Wall thickness/mm |
| --- | --- | --- |
| The number of vascular bundles | 0.907** | 0.877** |
| The distribution density of vascular bundles | -0.643** | -0.821** |
| The area of fiber sheath/mm<sup>2</sup> | 0.944** | 0.866** |
| Single fiber sheath area/mm<sup>2</sup> | 0.674** | 0.767** |
| Length/mm | 0.654** | 0.835** |
| Width/mm | 0.750** | 0.781** |

\*\*. Significantly correlated at the 0.01 level (bilateral)

Figure 2 shows the relationship between the outer circumference of the bamboo ring and the characteristics of vascular bundle. Figure 3 shows the relationship between the wall thickness of the bamboo ring and the characteristics of vascular bundle. The above six groups of data were analyzed by univariate linear regression to judge whether it belongs to the linear relationship. The goodness of fit (R^2^), significance test of a regression equation (F value), and significance test of the regression coefficient (P value) are shown in Table 3. The F value and P value were both significant. The characteristics of vascular bundle has a good linear correlation with the outer circumference and the wall thickness of bamboo.

**Table 3.** Regression analysis between the macro characteristics and the vascular bundle information

| Independent variable | Dependent variable | Regression equation | R<sup>2</sup> | F value | P value | Significance |
| --- | --- | --- | --- | --- | --- | --- |
| Outer circumference | The number of vascular bundles | $y=19.60x+42.11$ | 0.8229 | 125.47 | 1.18E-11 | significant |
| | The distribution density of vascular bundles | $y=-0.022x+7.230$ | 0.4140 | 19.07 | 1.66E-4 | significant |
| | The area of fiber sheath/mm<sup>2</sup> | $y=1.81x - 64.80$ | 0.8915 | 221.94 | 1.52E-14 | significant |
| | Single fiber sheath area/mm<sup>2</sup> | $y=0.0004x+0.0221$ | 0.4541 | 22.46 | 6.13E-5 | significant |
| | Length/mm | $y=0.0013x+0.2777$ | 0.4279 | 20.19 | 1.18E-4 | significant |
| | Width/mm | $y=0.0012x+0.2400$ | 0.5618 | 34.61 | 2.88E-6 | significant |
| Wall thickness | The number of vascular bundles | $y=545.32x - 131.38$ | 0.7694 | 80.09 | 4.10E-9 | significant |
| | The distribution density of vascular bundles | $y=-0.77x+8.34$ | 0.6746 | 49.74 | 2.72E-7 | significant |
| | The area of fiber sheath/mm<sup>2</sup> | $y=50.89x - 90.98$ | 0.7504 | 75.16 | 5.26E-9 | significant |
| | Single fiber sheath area/mm<sup>2</sup> | $y=0.0109x+0.0175$ | 0.5884 | 38.59 | 1.21E-6 | significant |
| | Length/mm | $y=0.0383x - 0.2425$ | 0.6973 | 62.21 | 1.77E-8 | significant |
| | Width/mm | $y=0.0271x+0.2397$ | 0.6106 | 42.33 | 5.63E-7 | significant |

a=0.05

**Figure 2.**
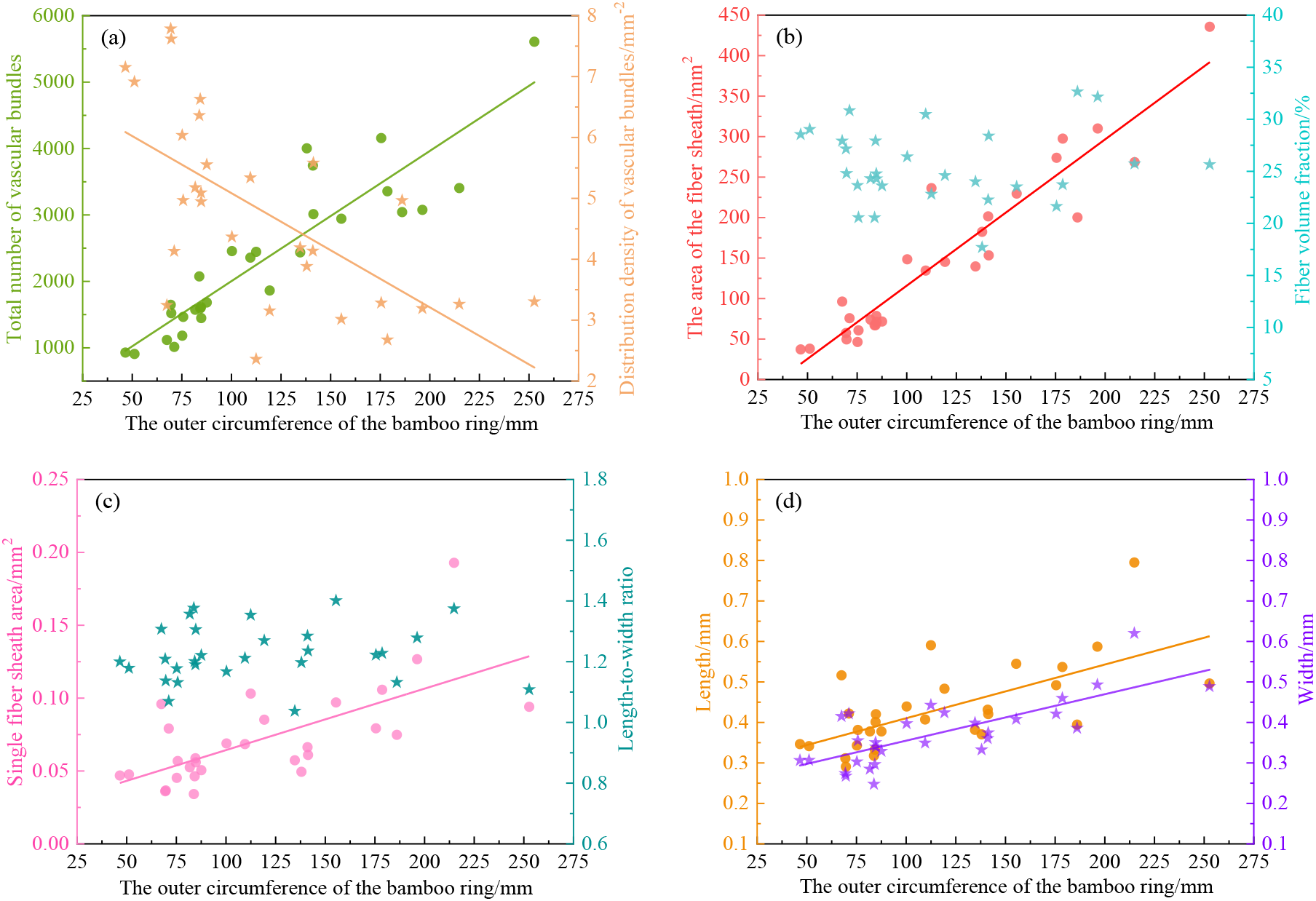
Relationship between the outer circumference of the bamboo ring and the characteristics of vascular bundle: **a** the total number of vascular bundles and the distribution density of vascular bundles; **b** the total area of fiber sheath and the fiber volume fraction; **c** the mean value of single fiber sheath area and the mean value of the length-to-width ratio of vascular bundle; **d** the mean value of the length of vascular bundle and the mean value of the width of the vascular bundle

**Figure 3.**
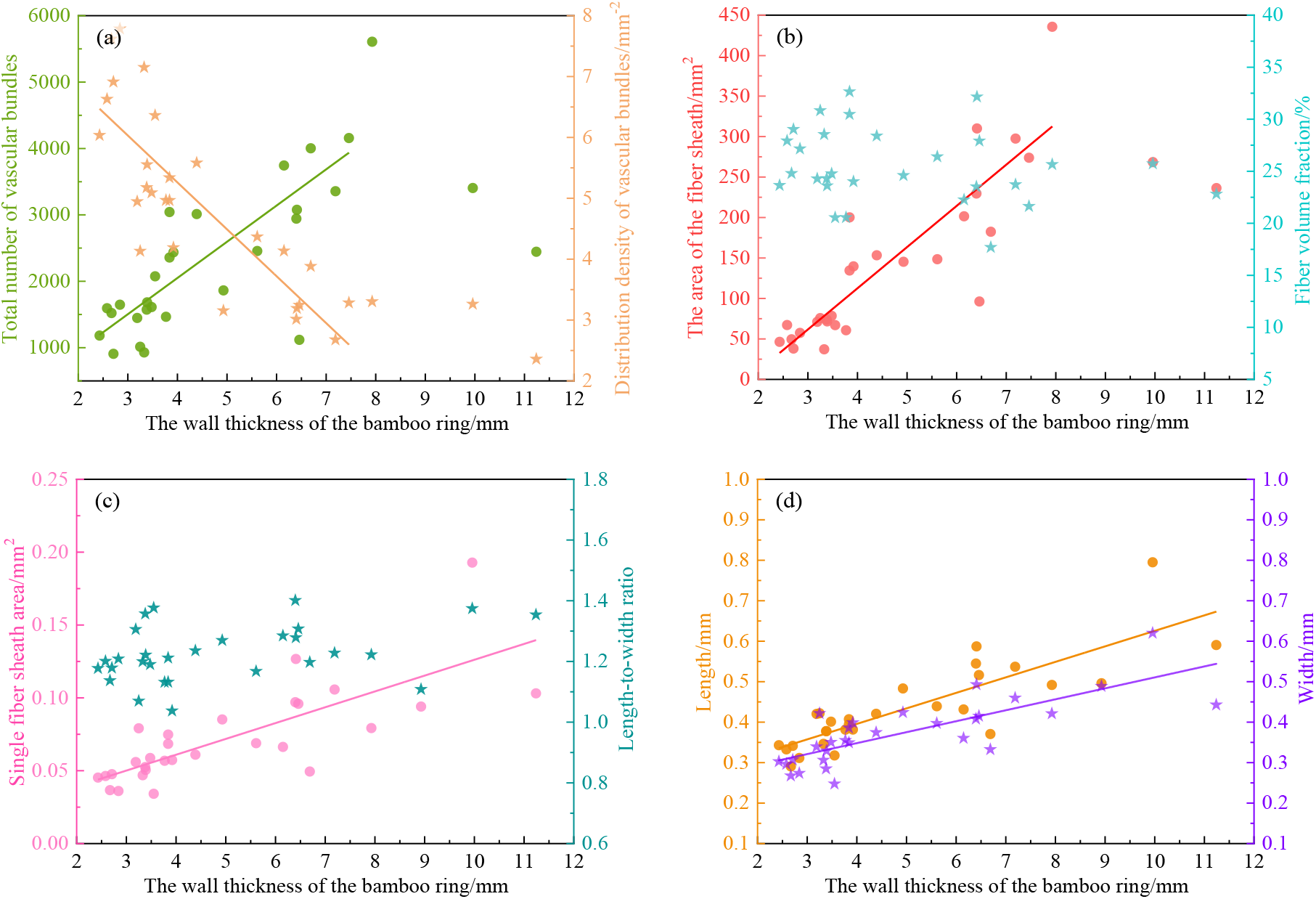
Relationship between the wall thickness of the bamboo ring and the characteristics of vascular bundle: **a** the total number of vascular bundles and the distribution density of vascular bundles; **b** the total area of fiber sheath and the fiber volume fraction; **c** the mean value of single fiber sheath area and the mean value of the length-to-width ratio of vascular bundle; **d** the mean value of the length of vascular bundle and the mean value of the width of the vascular bundle

### 2.3 The vascular bundle characteristics along the radial direction

#### 2.3.1 Fiber volume fraction

Referring to Table 5, the cross sections of bamboo were divided into layers according to their wall thickness, and the fiber volume fraction of each layer were obtained. A non-linear curve was fitted to the radial distribution of fiber volume fraction by the ExpDec1 function, and the evaluation indicators of the fitting results: the goodness of fit (R^2^) and the Reduced Chi-sqr are shown in Figure 4. The mean value of R^2^ was as high as 0.956, with the coefficient of variation was only 0.027 and the Reduced Chi-sqr are all less than 20, with a mean value of 7.757. This indicates that the ExpDec1 function can provide a good fit for the radial gradient structure of twenty-nine kinds of *Phyllostachys*.

**Figure 4.**
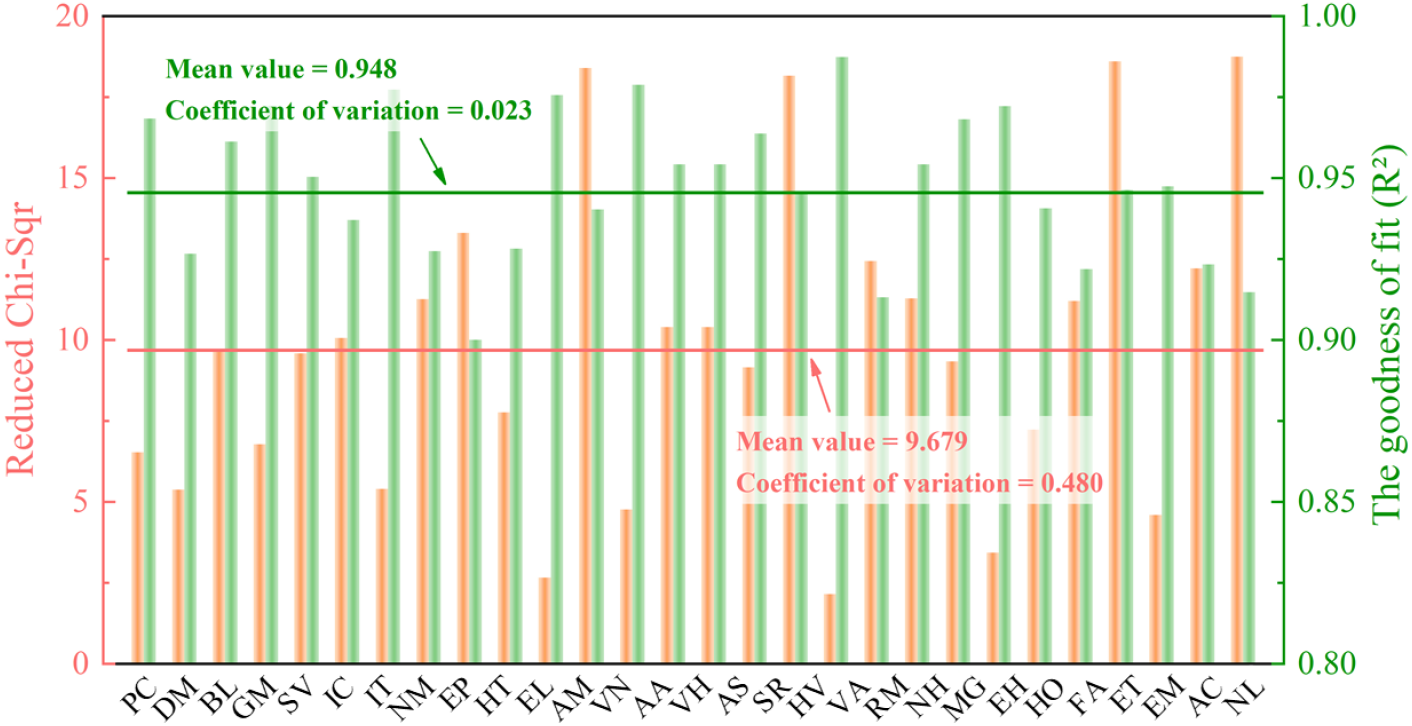
The fitting effect of ExpDec1 function of radial distribution of fiber volume fraction

The parameters <u>***A***</u> and <u>***t***</u> of twenty-nine kinds of *Phyllostachys* can be seen in Figure 5, which have an important influence on the ExpDec1 function. Although the radial distribution of fiber volume fraction of *Phyllostachys* all decreased exponentially, but the parameters <u>***A***</u> and <u>***t***</u> of different kinds of *Phyllostachys* do not correlate well with each other. The mean value of parameter A was 57.44 and the coefficient of variation was 0.27; the mean value of parameter <u>***t***</u> was 0.53, but the coefficient of variation was as high as 0.62, which was not very satisfactory. In this regard, we believed that mainly due to the adjustments made by different bamboo species in response to external environmental loads and internal physiological stresses. For bamboo species with higher axial height and thicker wall thickness, they are subjected to higher self-loads and higher natural stresses from external forces such as wind, rain, and snow are blowing. Therefore, it was necessary to optimize the distribution of vascular bundles to ensure its survival.

**Figure 5.**
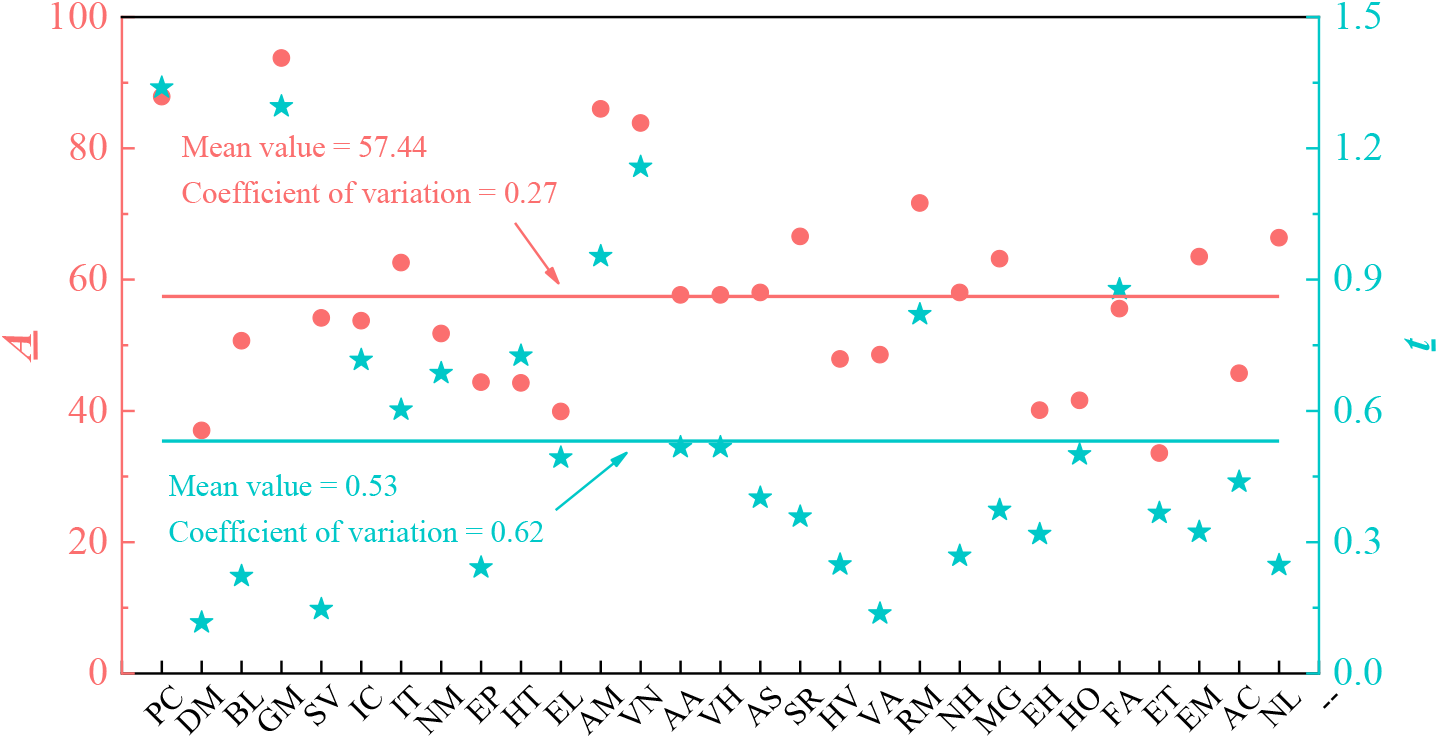
Mean value and coefficient of variation of <u>***A***</u> and <u>***t***</u> of ExpDec1 function

#### 2.3.2 The shape of vascular bundle

From the outer skin to the inner skin of bamboo, the shape of vascular bundles has a certain gradient variability. The vascular bundles on the outer skin are smaller in shape and densely distributed, whereas the vascular bundles on the inner skin are larger in shape and sparsely distributed. The different types of vascular bundles in the cross-section of *Phyllostachys* was shown in Figure 6. The vascular bundles of the semi-differentiated type have conducting tissue and they are arranged closely and staggeringly. The semi-open vascular bundles have obvious vessels and sieve tube, and their fiber sheath are divided into two valves, the lateral fiber sheath and the inner fiber sheath connected with each other. The vascular bundles of the open type appear in the middle and inside of the bamboo, the four fiber sheaths are independent, and the vessel, sieve tube and phloem are fully grained, representing the characteristic shape of specific bamboo species (Wen & Zhou, 1984; Wen & Zhou, 1985; Shang et al., 2015).

**Figure 6.**
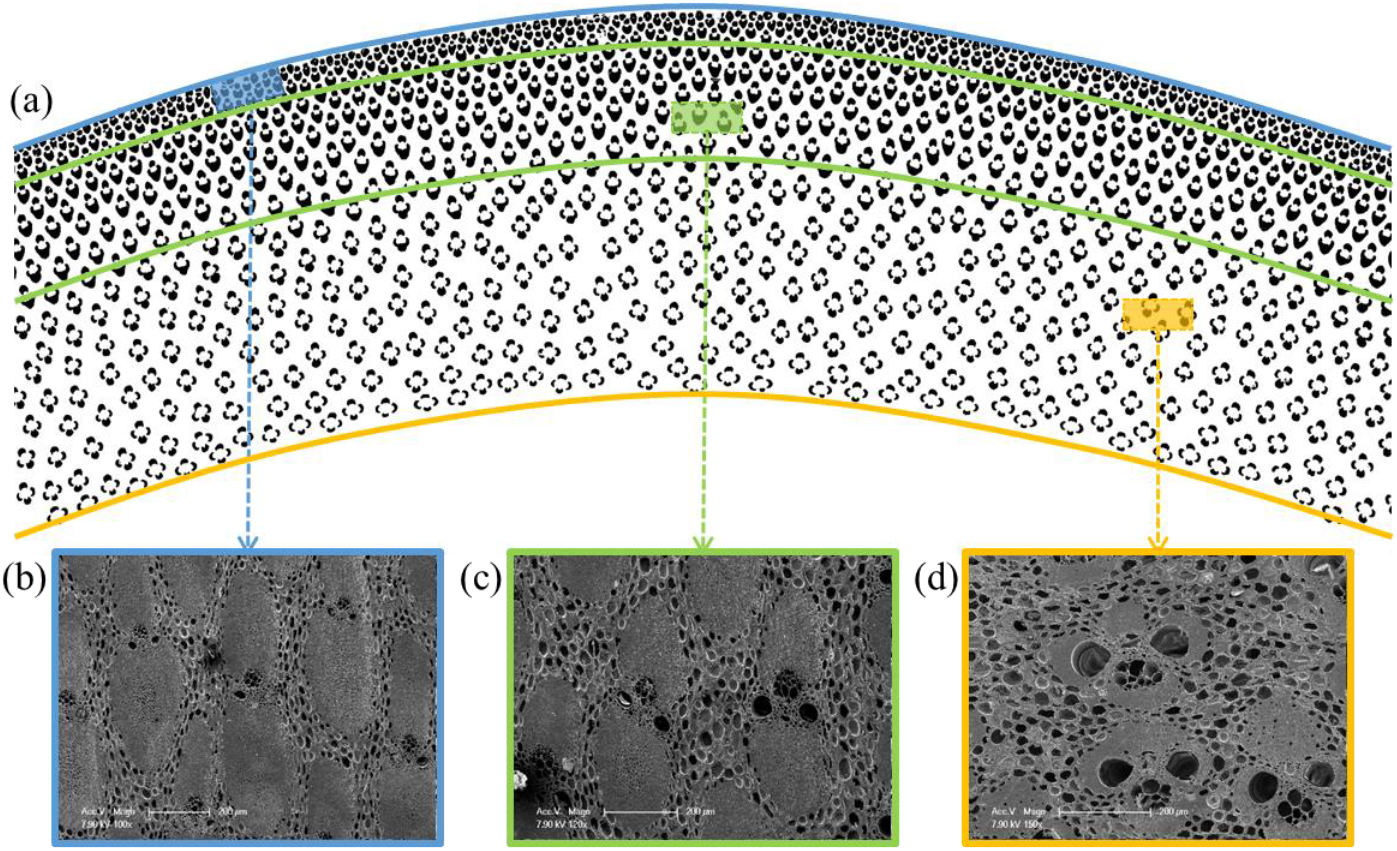
**a** Three types of vascular bundles in bamboo cross section; **b** The semi-differentiated type vascular bundle; **c** The semi-open type vascular bundle; **d** The open type vascular bundle

Along the radial direction, the changes of the shape of vascular bundle is shown in Figure 7. At the outer skin of bamboo, the length of the vascular bundle was longer than the width of the vascular bundle and the length-to-width ratio of which was about 1.5. When reaching the position 30-40% away from the outer skin, the length-to-width ratio of the vascular bundle of some bamboo species increased and the others were consistent with that of the outer skin, but the length and width of the vascular bundle were larger than that of the outer skin. When in the radial middle position, the length of the vascular bundle reaches the maximum, and the length-to-width ratio of the vascular bundle begins to decrease gradually. When reaching the position 70-80% away from the outer skin, the length-to-width ratio of the vascular bundle was about 1:1, and the length of the vascular bundle remains unchanged or decreases slightly. At the inner skin of bamboo, the width of the vascular bundle reached the maximum and the length-to-width ratio of the vascular bundle was about 0.75. Besides, the Quadratic function and linear function were used to fit the radial distribution of the length-to-width ratio and the width of the vascular bundle. The goodness of fit (R^2^) was shown in Figure 8, the average value of R^2^ was as high as 0.869 and 0.926, and the coefficient of variation was only 0.089 and 0.059. It shows that the shape of vascular bundle also had a high degree of consistency in the radial direction.

**Figure 7.**
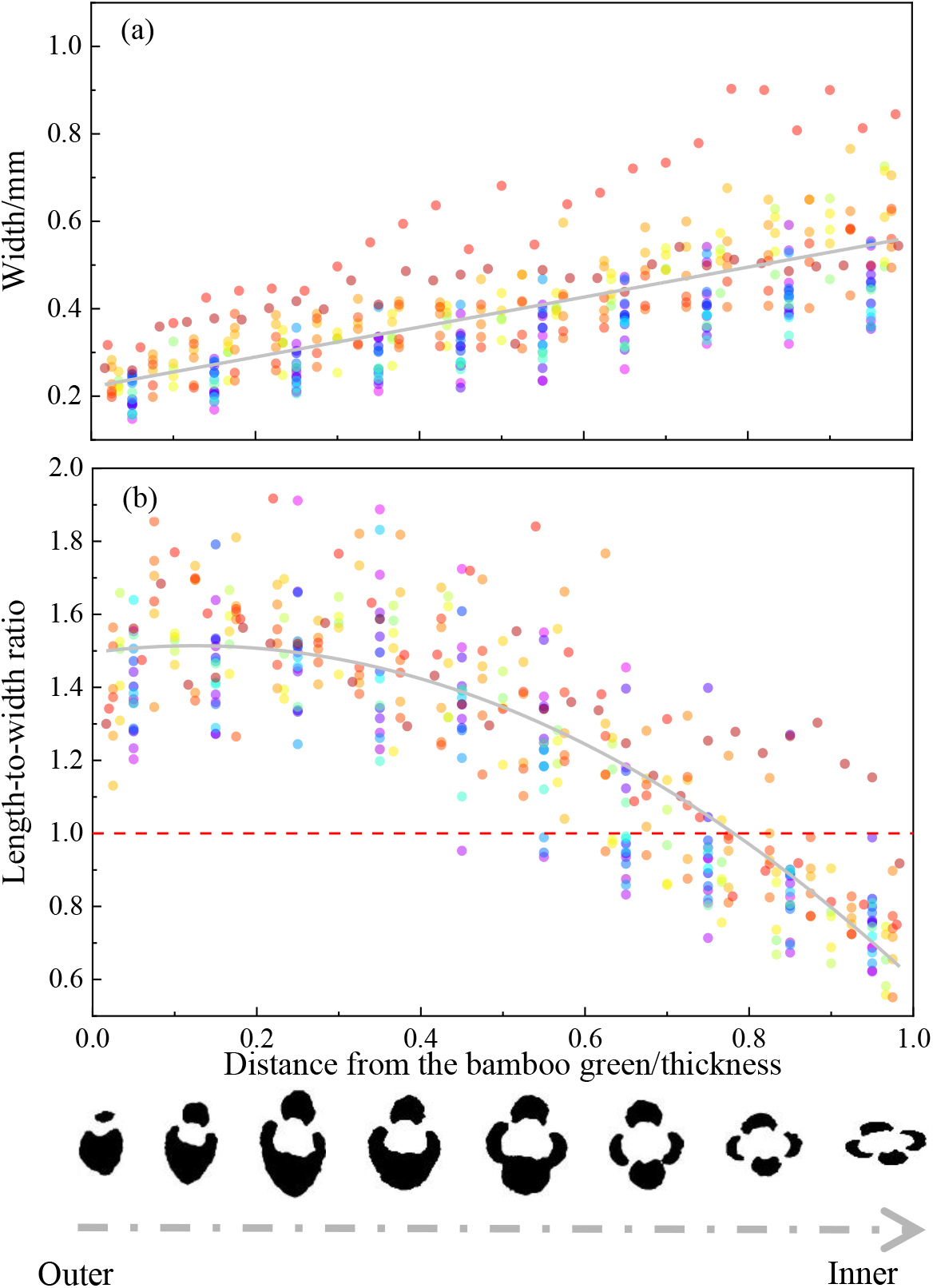
The shape of vascular bundle changed along the radial direction: **a** change in width; **b** change in the length-to-width ratio

**Figure 8.**
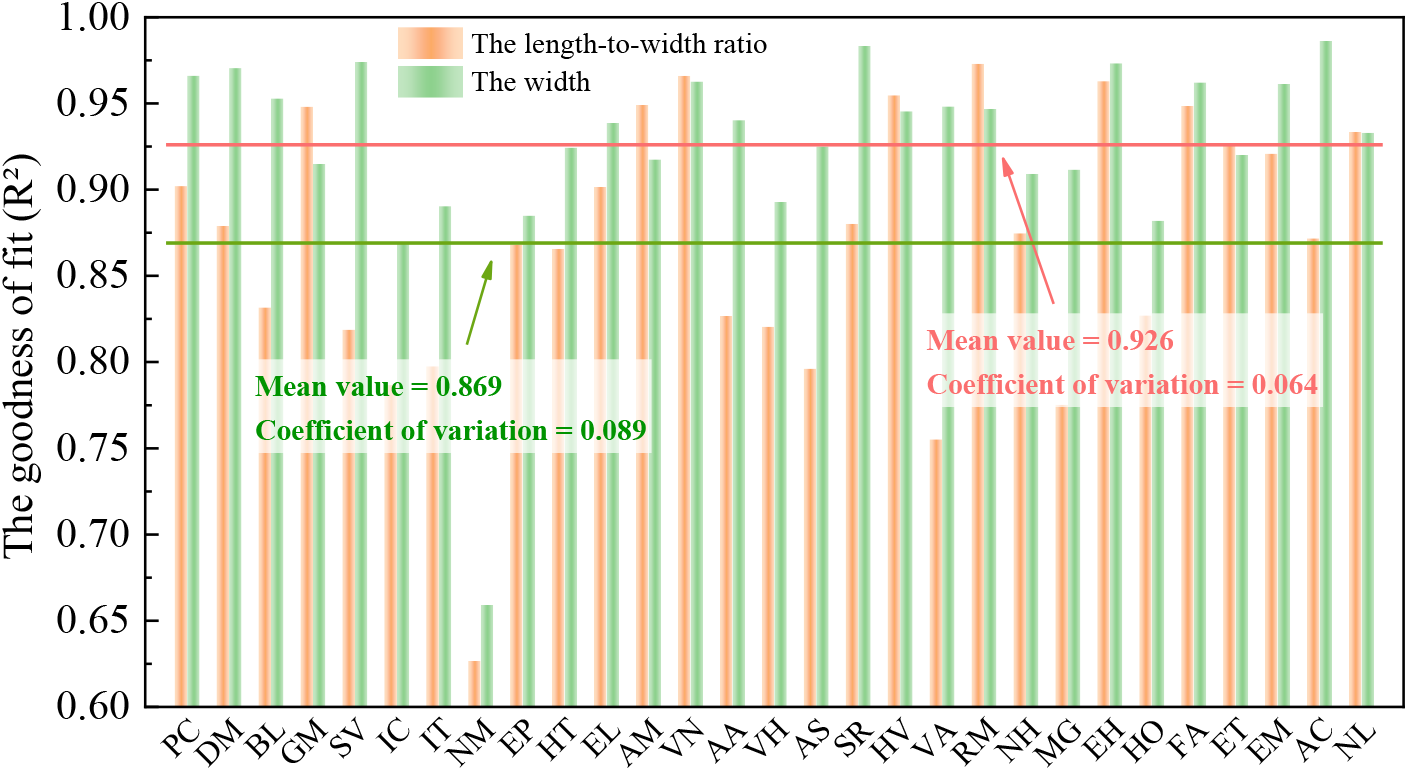
The fitting effect of radial distribution of the length-to-width ratio and the width of vascular bundle

## 3. Conclusion

This study constructed a universal detection model for vascular bundle, which can automatically and accurately analyze the characteristics of vascular bundles for *Phyllostachys*. The precision rate of the model was 96.97%. The characteristics of vascular bundle were revealed and the conclusions were as follows: 1) The total number of vascular bundles, the distribution density of vascular bundles, the total area of fiber sheath, the mean value of single fiber sheath area, the mean value of the length of the vascular bundles and the mean value of the width of vascular bundles had a positive and negative linear correlation with the outer circumference and the wall thickness of bamboo, respectively. 2) The difference of fiber volume fraction and the mean value of the length-to-width ratio of vascular bundle between twenty-nine bamboo species were relatively small. The average fiber volume fraction was 25.50 ± 3.51% and the average of the mean value of the length-to-width ratio of the vascular bundle was 1.226 ± 0.091. 3) The radial distribution of fiber volume fraction decreased exponentially, and the R^2^ was 0.948. The radial distribution of the length-to-width ratio of the vascular bundle decreased quadratically and the R^2^ was 0.869. The radial distribution of the width of the vascular bundle increased linearly and the R^2^ was 0.926.

## 4. Materials and methods

### 4.1 Preparing samples

Twenty-nine kinds of three-year-old *Phyllostachys* species were collected from China (Table 4). Bamboo rings with 20 mm high were cut from the 2-meter height above the ground of each bamboo culm. Bamboo rings (free from defects) were conditioned at 20°C and 60% relative humidity (RH) for 28 days in a temperature humidity chamber (HWS-250, SAIFU, China). Sanding pads with 320 mesh (NO28976, PROXXON, Germany) were used during the sanding process. The pores in vessels and phloems were filled during the sanding process, which can provide an improvement in the accuracy of detecting the fiber sheath. Bamboo rings were scanned at the cross-section by a scanner (PERFECTION V850 PRO, EPSON, Japan) in 16-gray mode with a resolution of 9600 ppi (Figure 9). The fiber sheaths and parenchyma cells were observed in the scanned images clearly.

**Table 4.** Macro characteristics of *Phyllostachys*

| No. | Species | Wall thickness / mm | Outer circumference / mm |
| --- | --- | --- | --- |
| PC | *Phyllostachys parvifolia* C. D. Chu & H. Y. Chou | 3.84 | 186.09 |
| DM | *Phyllostachys dulcis* McClure | 3.55 | 83.83 |
| BL | *Phyllostachys bambusoides* f. *lacrima-deae* | 3.39 | 87.58 |
| GM | *Phyllostachys glauca* McClure | 5.61 | 100.27 |
| SV | *Phyllostachys sulphurea* var. *viridis* R. A. Young | 3.92 | 134.69 |
| IC | *Phyllostachys iridescens* C. Y. Yao & S. Y. Chen | 2.84 | 69.41 |
| IT | *Phyllostachys incarnata* T. W. Wen | 3.84 | 109.59 |
| NM | *Phyllostachys nidularia* Munro | 3.38 | 81.75 |
| EP | *phyllostachys edulis* cv. *Pachyloen* | 11.24 | 112.46 |
| HT | *Phyllostachys heterocyclus* cv. *Tao Kiang* | 6.41 | 196.29 |
| EL | *Phyllostachys edulis* cv. *Luteosulcata* | 7.19 | 178.66 |
| AM | *Phyllostachys aureosulcata* McClure | 2.43 | 75.20 |
| VN | *Phyllostachys vivax* McClure f. *aureocaulis* N.X. Ma | 4.93 | 119.29 |
| AA | *Phyllostachys aureosulcata* McClure *Aureocaulis* | 3.48 | 84.65 |
| VH | *Phyllostachys vivax* McClure cv. *Huanwenzhu* | 6.4 | 155.42 |
| AS | *Phyllostachys aureosulcata* cv. *Spectabilis* | 3.19 | 84.78 |
| SR | *Phyllostachys sulphurea* (Carr.) Rivière & C. Rivière | 9.96 | 214.87 |
| HV | *Phyllostachys heterocyclus* cv. *Viridisulcata* | 6.69 | 138.00 |
| VA | *Phyllostachys viridi-glaucescens* (Carr.) A. et C. Riv. | 3.77 | 75.71 |
| RM | *Phyllostachys rubromarginata* McClure | 3.25 | 71.20 |
| NH | *Phyllostachys nigra* var. *henonis* Mitford Stapf ex Rendle | 4.39 | 141.34 |
| MG | *Phyllostachys edulis* (Carr.) H. De Lehaie | 8.93 | 271.66 |
| EH | *Phyllostachys mannii* Gamble | 3.33 | 46.55 |
| HO | *Phyllostachys heterocyclus* cv. *Obliquinoda* | 7.46 | 175.54 |
| FA | *Phyllostachys flexuosa* (Carr.) A. et C. Riv. | 2.58 | 84.27 |
| ET | *Phyllostachys edulis* f. *Tubaeformis* | 6.15 | 141.06 |
| EM | *Phyllostachys elegans* McClure | 6.46 | 67.46 |
| AC | *Phyllostachys atrovaginata* C. S. Chao & H. Y. Chou | 2.67 | 69.66 |
| NL | *Phyllostachys nigra* Lodd. ex Lindl. Munro | 2.71 | 51.11 |

**Figure 9.**
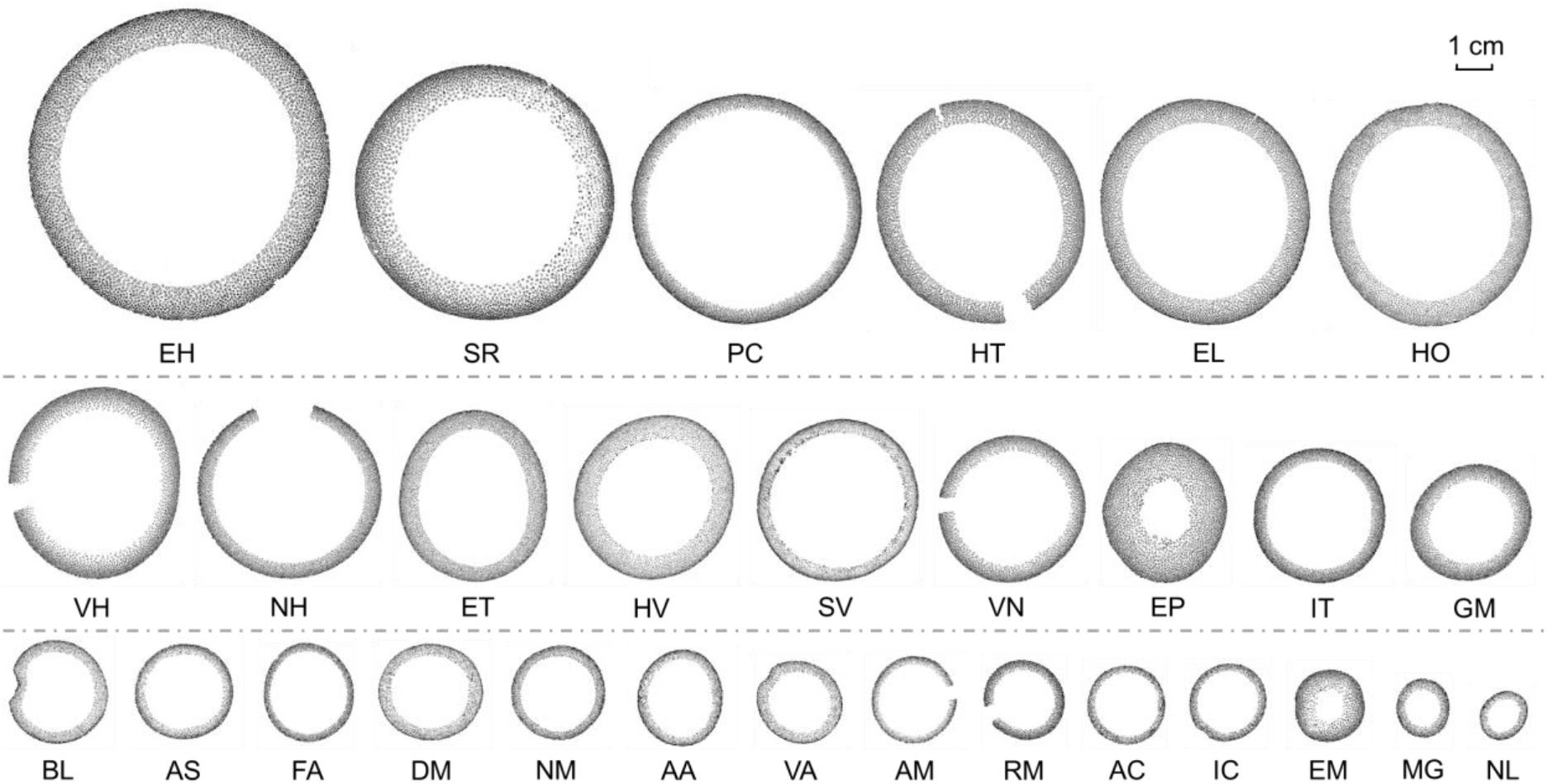
The binarization of the cross section of bamboo rings

### 4.2 Constructing a detection model for vascular bundles of Phyllostachys

Figure 10 shows the process of the model construction. The original scanned cross-sectional images of twenty-nine kinds of *Phyllostachys* were divided into several smaller patches of 1024 × 1024 pixels, and the vascular bundles in the images were labelled. Two thousand vascular bundles were labelled from each kind of bamboo species as a training data set. Due to the limited area of the vascular bundles on the outer skin of bamboo, it was a small target in the whole image and was easily ignored in the actual detection process, eventually resulting in inaccurate detection. Therefore, we increased the number of this kind of vascular bundle in the labeling process. The suggested labeling ratio was 6:4 of vascular bundles on the outer skin and inner skin. The labelled vascular bundles of each bamboo species were mixed together and trained by the YOLO (You Only Look Once, a stable and popular machine learning method) algorithm. **[Figure 10]**

**Figure 10.**
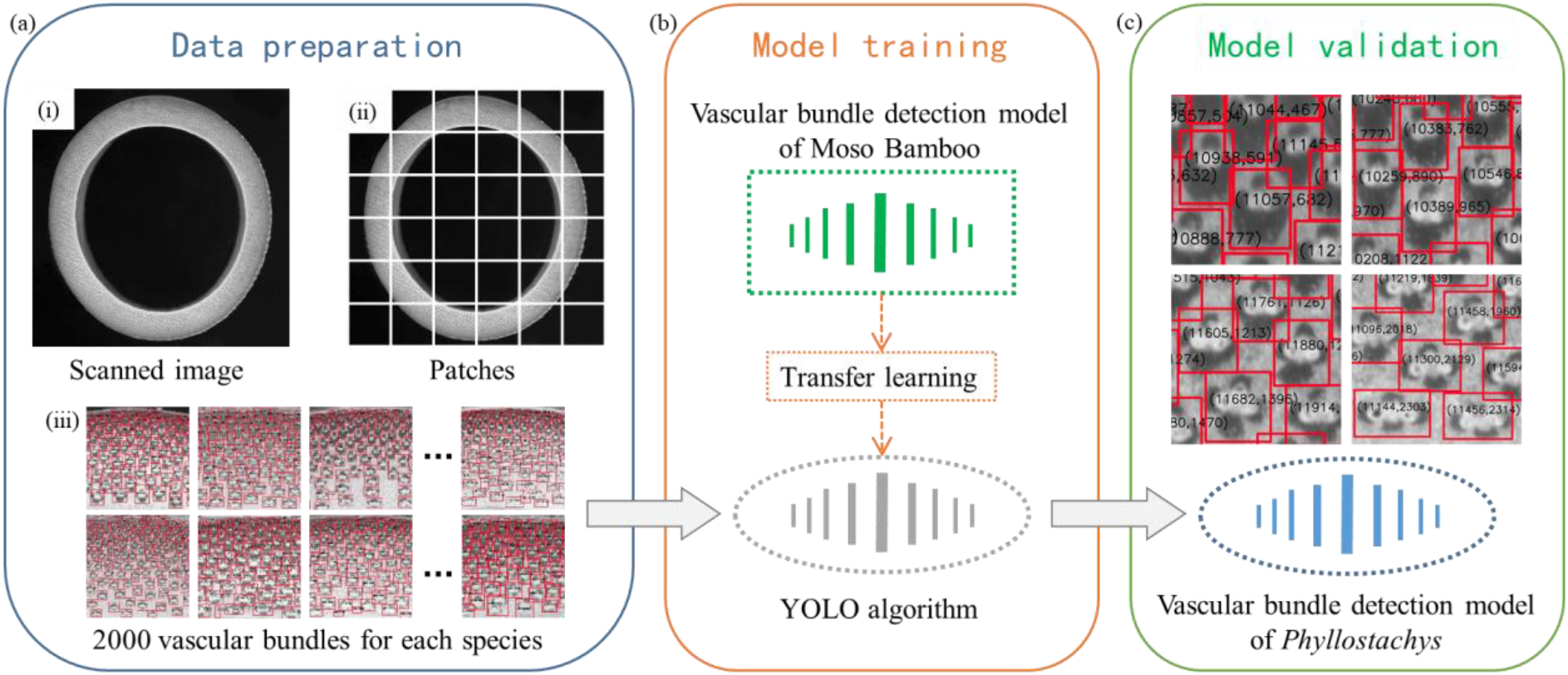
The process of model construction: **a** data preparation; **b** model training; **c** model validation

In model training, since the sample size of the current data set was relatively small (2000 vascular bundles for each species, only 1/10 of the labels when training the “vascular bundle detection model” of Moso bamboo), and the difference in vascular bundles between *Phyllostachys* was not significant, so the transfer learning was used in this research. Based on a large number of vascular bundle features that have been learned by the “vascular bundle detection model” of Moso bamboo, these features are applied to this model training through transfer learning, saving a lot of computing power and time of Graphics Processing Unit (GPU) for Feature Engineering. Finally, we applied the trained model to the original scanned image to verify the accuracy of the model. For bamboo species with poor detection results, the error can be reduced by increasing the number of their labeled vascular bundles.

### 4.3 Acquiring vascular bundle characteristics in cross section

The universal vascular bundle detection model was used to analyze twenty-nine kinds of *Phyllostachys* based on their cross section images, the number of vascular bundles and the distribution density of vascular bundle can be obtained directly. Based on the identification of vascular bundles by the detection model, we used K-Means clustering algorithm (a machine learning method that automatically clusters samples based on defined characteristics), which cluster the pixel points with large pixel value into white, and the pixel points with small pixel value into black to realize the binary of vascular bundles. Then we multiplied the total number of black pixel points in the overall image by the actual area of each pixel (6.9696 μm^2^, represented by corresponding resolution of 9600 ppi) can obtain the total area of fiber sheath. The fiber volume fraction can be calculated by dividing the cross-sectional area of fiber sheath by the cross-sectional area of bamboo culm. Furthermore, every vascular bundles in the cross section were selected, and the single fiber sheath area, the length-to-width ratio, the length and the width of each vascular bundle were obtained by the vascular bundle detection model, and their mean values were calculated.

### 4.4 Acquiring vascular bundle characteristics along the radial direction

#### 4.4.1 Fiber volume fraction

According to the comparison table between the bamboo wall thickness and the number of layers reported by Xu et al. (2021), we divided the binarized cross section images into layers corresponding to their wall thickness along the radial direction uniformly (Table 5).

**Table 5.** Comparison between the number of layers and the wall thickness

| Wall thickness / mm | 2~4 | 4~6 | 6~8 | 8~10 | 10~12 |
| --- | --- | --- | --- | --- | --- |
| Number of layers | 10 | 15 | 20 | 25 | 30 |

In this research, the fiber volume fraction (y) was set to vary continuously in one dimension along the radial direction (x-direction) of bamboo, and nonlinear curve fitting was performed on the scatter plot of the radial distribution of fiber volume fraction based on the decay exponential function (ExpDec1 function, Eq.1).

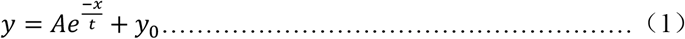

Where <u>***A***</u> is the constant term of the function. If <u>***A***</u> is larger, to some extent the total fiber volume fraction of that bamboo species is larger. <u>***e***</u> is the base of the natural logarithm. <u>***t***</u> is the constant term of the exponent <u>***x***</u>, which directly affects the final two-dimensional shape of the function. <u>***y***</u> can be considered as a correction. Therefore, <u>***A***</u> and <u>***t***</u> are of significance for the final expression form of the exponential function. In this research, the parameters <u>***A***</u> and <u>***t***</u> obtained from the fitting for the radial distribution of fiber volume fraction of twenty-nine kinds of *Phyllostachys* were counted and compared.

#### 4.4.2 The shape of vascular bundle

In order to express the changes in the shape of the vascular bundle in the radial direction, the length-to-width ratio, the length and the width of the vascular bundle were used as indexes. According to the above-mentioned number of layers for every bamboo species, fifteen vascular bundles were selected in each layer, and calculates the mean value of its shape parameters (Figure 11).

**Figure 11.**
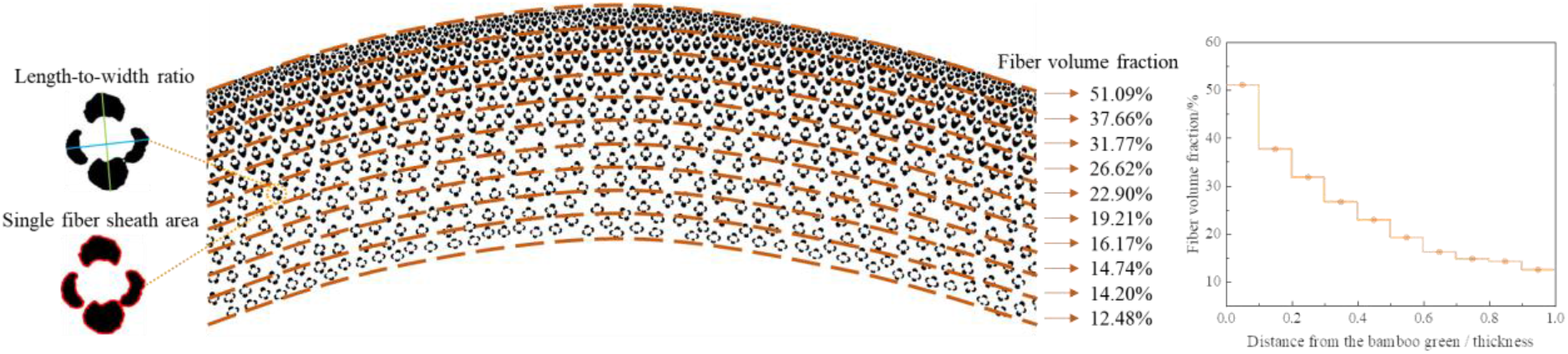
Acquire the characteristics of vascular bundle in radial direction

## Acknowledgements

The authors would like to thank Professor Zehui Jiang and her team at the International Center for Bamboo and Rattan for experimental venues and equipment provided. This work was supported by the Basic Scientific Research Funds of the International Center for Bamboo and Rattan (Grant No. 1632020012) and the National Natural Science Foundation (Grant No. 32071855).

